# Tapetum specific TA29 promoter is regulated by *cis*- elements binding to positive and negative regulators

**DOI:** 10.1101/2024.12.27.630565

**Authors:** Preeti Apurve Sharma, Kamlesh Kumar Soni, Bm Minhajuddin, Satish Kumar Yadava, Pradeep Kumar Burma

## Abstract

The TA29 gene from *Nicotiana tabacum* was one of the first tapetum-specific genes identified in 1990. Its promoter has been successfully used to develop *barnase*/*barstar* gene-based male sterility and restorer transgenic lines for hybrid seed production in crops such as mustard. Despite its biotechnological significance, no major studies have been conducted to delineate the regulatory mechanisms of its promoter. The presented work identifies critical *cis*-elements of the promoter, starting with *in-silico* analysis followed by functional validation through mutational studies and promoter activity analysis in transgenic tobacco lines. This work led to the identification of *cis*-elements that are putative binding sites for the transcription factors NtMYB12, NtGT-1a, NtNAG1, and NtLZ-HD. The binding sites for NtMYB12, NtGT-1a, and NtNAG1 act as positive regulators, whereas the binding site for NtLZ-HD functions as a negative regulator.

## Introduction

The tapetum-specific *TA29* gene from *Nicotiana tabacum* was discovered in 1990 by Robert Goldberg’s group (Koltunow et al. 1990, first report) while identifying specific gene sets for anther development. Several anther-specific genes, including *TA29*, were identified through differential screening of anther cDNA libraries, Northern hybridizations and other analyses. *In situ* hybridization showed that *TA29* expressed specifically in the tapetum. Deletion analysis of a ~5.6 Kb region upstream to the putative transcriptional start site, showed that a −279 bp long promoter fragment sufficed for tapetum-specific expression, with a 122 bp region spanning −207 and −85 being crucial.

Further, it was demonstrated that expression of genes like *barnase* from *Bacillus amyloliquefaciens*, under TA29 promoter, leads to male sterility (Mariani et al. 1990). Notably, the promoter also exhibited tapetum specific activity in other plants like oilseed rape *Brassica napus*. The identification of the TA29 promoter enabled the development of a pollination control system for hybrid seed production in various crops, using the *barnase* and *barstar* genes using e.g. the 863 bp promoter (Jagannath et al. 2001, 2002).

Despite their widespread application, research on the regulatory mechanisms of tapetum-specific promoters, including transcription factors and *cis*-regulatory modules, remains limited. Verma and Burma (2017) investigated regulation of a tapetum specific promoter from *Arabidopsis thaliana* and found it to be controlled by both positive (AtMYB1 and AtMYB80) and negative (AtMYB4) regulators. The positive regulators, AtMYB1 and AtMYB80, were observed to regulate the A9 promoter through specific *cis*-elements, ‘CACCAATCC’ at −57 bp and ‘CAACACCTC’ at −290 bp, respectively. On the other hand, the negative regulator AtMYB4, worked through three *cis*-elements viz., ‘TGGTTTGTT’ at −391 bp, ‘TGTAACCA’ at −350 bp, and ‘GTTGTGG’ at −291 bp.

Although both TA29 and A9 promoters show tapetum-specific activity in plants like tobacco, *Arabidopsis*, and *Brassica juncea*, the *cis*-elements in A9 are notably absent in TA29, indicating different regulatory mechanisms. This study aimed to identify the *cis*-elements regulating the TA29 promoter in tobacco.

## Materials and Methods

### Plant material and growth conditions

*Nicotiana tabacum* (tobacco) cv. Xanthi plants were used for the current study. Wild type and transgenic tobacco lines were either grown on MS medium in tissue culture room or on soil in green houses. In both the cases, plants were grown at 28^0^C under 16/8 h light to day cycle. Further, in the greenhouse humidity was maintained at ~ 70%.

### Development of constructs

A total of 9 different constructs were developed for this study. In each of the constructs the TA29 URM (wild type or mutated) were cloned upstream to a intron containing *β-glucuronidase* gene (Gi), (Vancanneyt et al. 1990) followed by a polyA (pA) signal of CaMV35S. The expression cassette was assembled in the vector pRT100 and finally cloned into the binary vector pZP200N (Bhullar et al. 2003). The identified *cis*-elements in the TA29 URM were mutated by replacing them random DNA stretches of same length. These were incorporated by PCR and the expression cassettes and binary vector assembled through homology based recombinational cloning (In-fusion HD Cloning Kit, Clonetech) or by standard restriction-ligation method. The constructs were confirmed by both restriction digestion and sequencing of various cassettes. The constructs are described in the results and discussion section.

### Development of transgenic lines

Tobacco leaf explants were used for Agrobacterium-mediated transformation following the protocol of Svab et al (1995). *Agrobacterium tumefaciens* strain GV2260 carrying different binary vectors were used for plant transformation. Following, co-cultivation of leaf explants they were placed on MS medium, supplemented with auxin (NAA, 0.1 mg/l), cytokinin (BAP,1.0 mg/l), kanamycin (100mg/ ml) and cefotaxime (200 mg/l). Regenerating shoots (one from each explant) from callus were transferred to MS media with kanamycin (100mg/ ml), for rooting. The transgenic plants were maintained in tissue culture on MS media supplemented with kanamycin (50 mg/ ml). For flowering, plants were transferred to soil and grown in green house.

### Measuring GUS activity in transgenic lines

Total protein from buds was extracted in GUS extraction buffer (Jefferson et al. 1987). Following estimation of protein concentration GUS activity was quantitated in the extract, using MUG (4-methylumberrifyl-β-glucuronide) as the substrate according to Jefferson et al. (1987). 1 and 2 μg of total protein was used in each case. Further, the assay was carried out for two time points e.g., 10 and 20 minutes. The product, methylumbelliferone (MU,) released was estimated in fluorimeter using excitation wavelength of 365 nm and emission recorded at 460 nm. GUS activity was expressed as picomoles of MU/min/mg total protein, using a standard curve of MU.

### Histochemical staining for GUS activity

Callus, stem and root tissues of transgenic lines developed with mLZ-HD construct was stained for GUS activity using 5-bromo-4-chloro-3-indolyl-β-D-glucuronic acid (X-gluc) in buffer as described in Jefferson et al (1987). Samples were incubated overnight at 37^0^C. After appropriate destaining they were observed under stereo-zoom microscope.

### Statistical analysis

JASP (version 0.18.3) was used for statistical analyses. Shapiro-Wilk test was carried out to test the normality of the data and GUS activity driven by different URMs compared using Welch’s two sample *t* test.

## Results and discussion

### Identifying putative *cis*-elements

To identify the putative *cis*-elements in the TA29 promoter, a 600 bp region upstream to the ‘translational start site’ (hitherto referred to as Upstream Regulatory Module, URM) for 24 genes reported to express in tapetum/anther tissue were analysed. Using a combination of MEME and MAST tools of MEME Suite 4.9.1, followed by manual curation, short sequence patterns across the 24 sequences were identified and consensus motifs were generated. These motifs were further analysed using PLACE and TRANSFAC for similarity with known *cis*-elements or transcription factor binding sites. As shown in Fig. 1a and detailed in Supplementary Fig. 1, this led to the identification of putative *cis*-elements binding to transcription factors (TFs) NtMYB12, NtGT-1a, NtNAG1, and NtLZ-HD in the TA29 URM. There are two sites each for NtMYB12 and NtLZ-HD located in the region between −200 and −217 for the former and at −786 and −136 for the latter. The single *cis*-element for NtNAG1 is located at −200 position. There are six putative *cis*-elements for NtGT-1a scattered at different locations across the URM including one site in the 5’UTR. The sequences of the identified *cis*-elements is presented in Fig. 1b.

**Fig. 1a.**
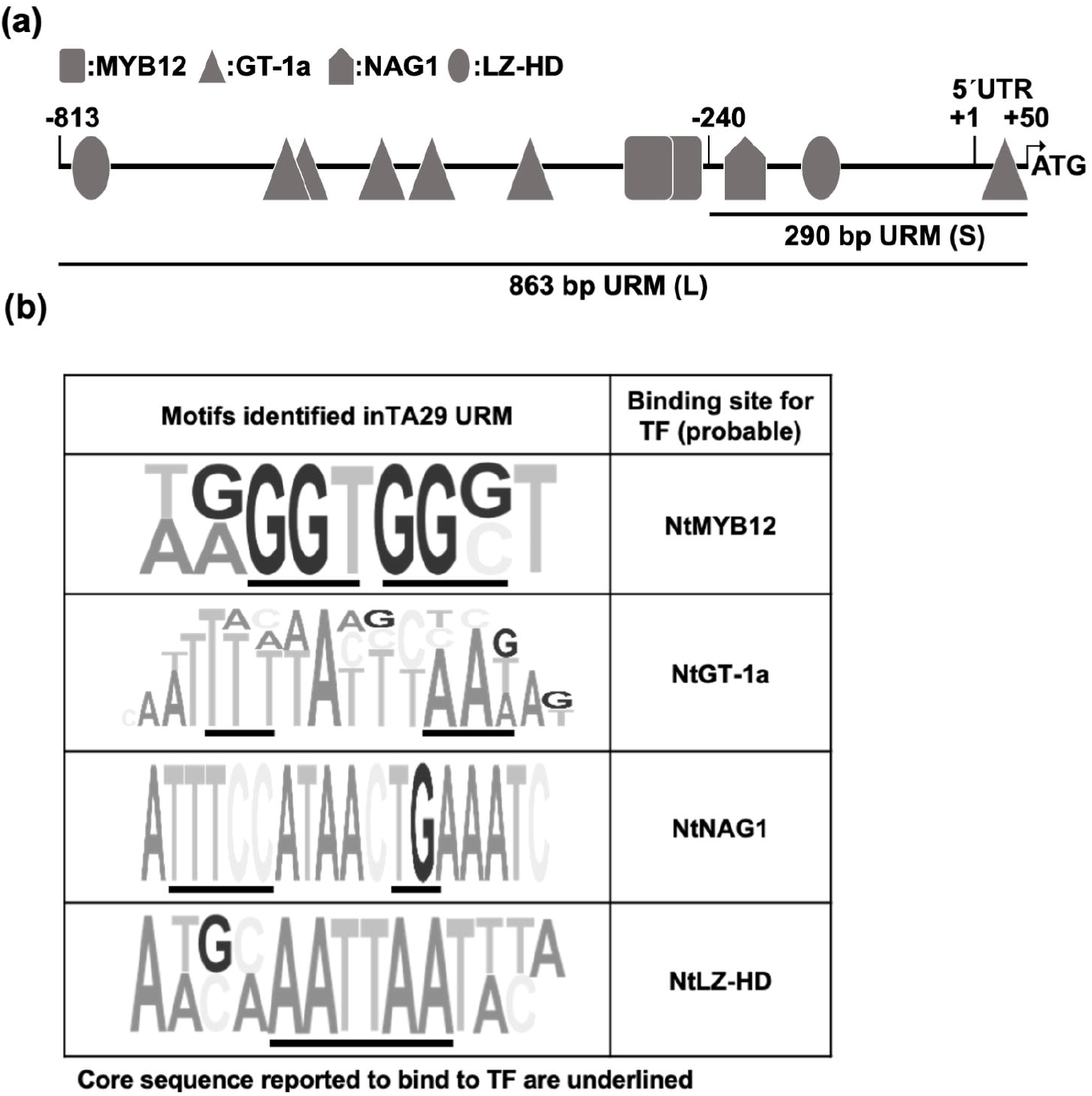
Organization of TA29 URM indicating the identified *cis*-elements. **b**. The identified motifs of *cis*-elements presenting sequence variation and transcription factor (TF) that probably bind to it. Core bases recognised by TF is underlined.

### Analysing the role of the identified cis-elements in URM activity

For experimentally testing the role of the identified *cis*-elements in regulation of the TA29 URM, different URM constructs encompassing mutated putative *cis*-elements for each TF were developed by replacing most of the key bases of the given region. These mutations were carried out in the background of either 863bp (L) or 290bp (S) long URM. This includes 50 bp of 5’UTR. The shorter version was designed with its distal end encompassing the 122 bp crucial region identified in the first report. An intron-containing *β-glucuronidase* gene (Gi) was cloned downstream of the URMs and binary vectors were developed for plant transformations (Fig. 2). The activity of the mutated URMs was analysed in transgenic tobacco lines and compared with the wild-type TA29 URMs of 863 and 290 bp lengths. Around 10 independent lines were analysed in each case. Activity was measured at 3 different bud stages, viz., BS1 (bud size of ~1 to 5 mm), BS2 (~6 to 10 mm), and BS3 (~11 to 15 mm). Several buds in the given size ranges were taken for each analysis. The tapetum layer is first seen in BS1, fully developed in BS2, and starts to degenerate by BS3. In the case of 290bp URM, studies were carried out in BS1 and BS2.

**Fig. 2.**
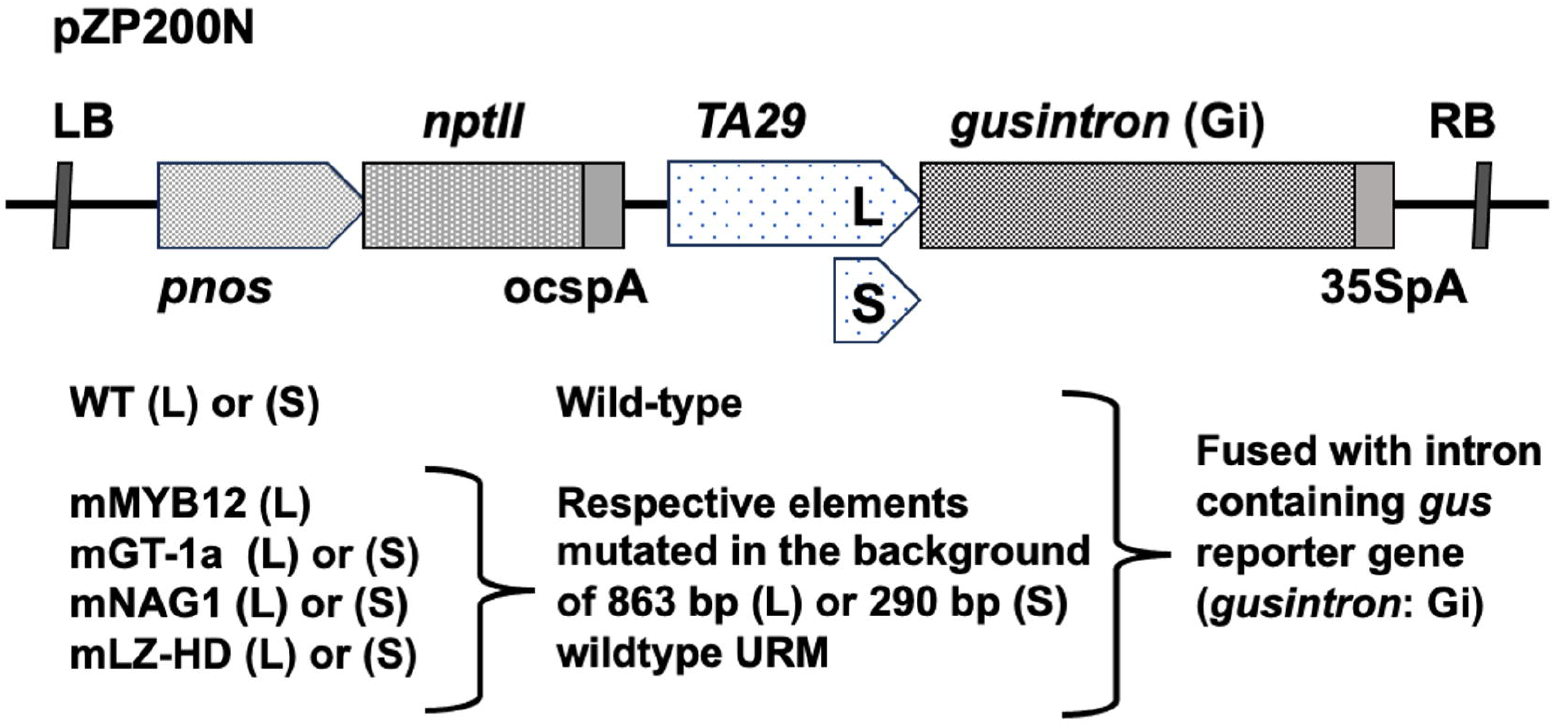
Outline of binary vectors used for transformation of tobacco.

The activity of the URMs was measured as GUS activity in each independent line (see Fig 3). The mean (± S.E.) GUS activities observed at BS1 and BS2 stages are summarized in Table 1 (for details see Supplementary Table 1). Shapiro-Wilk test for normality showed that the data for different transgenic lines for each construct showed normal distribution. Mean GUS activity obtained from mutant URMs were compared to their respective wild-type URM using Welch’s *t*-test.

**Table 1:**
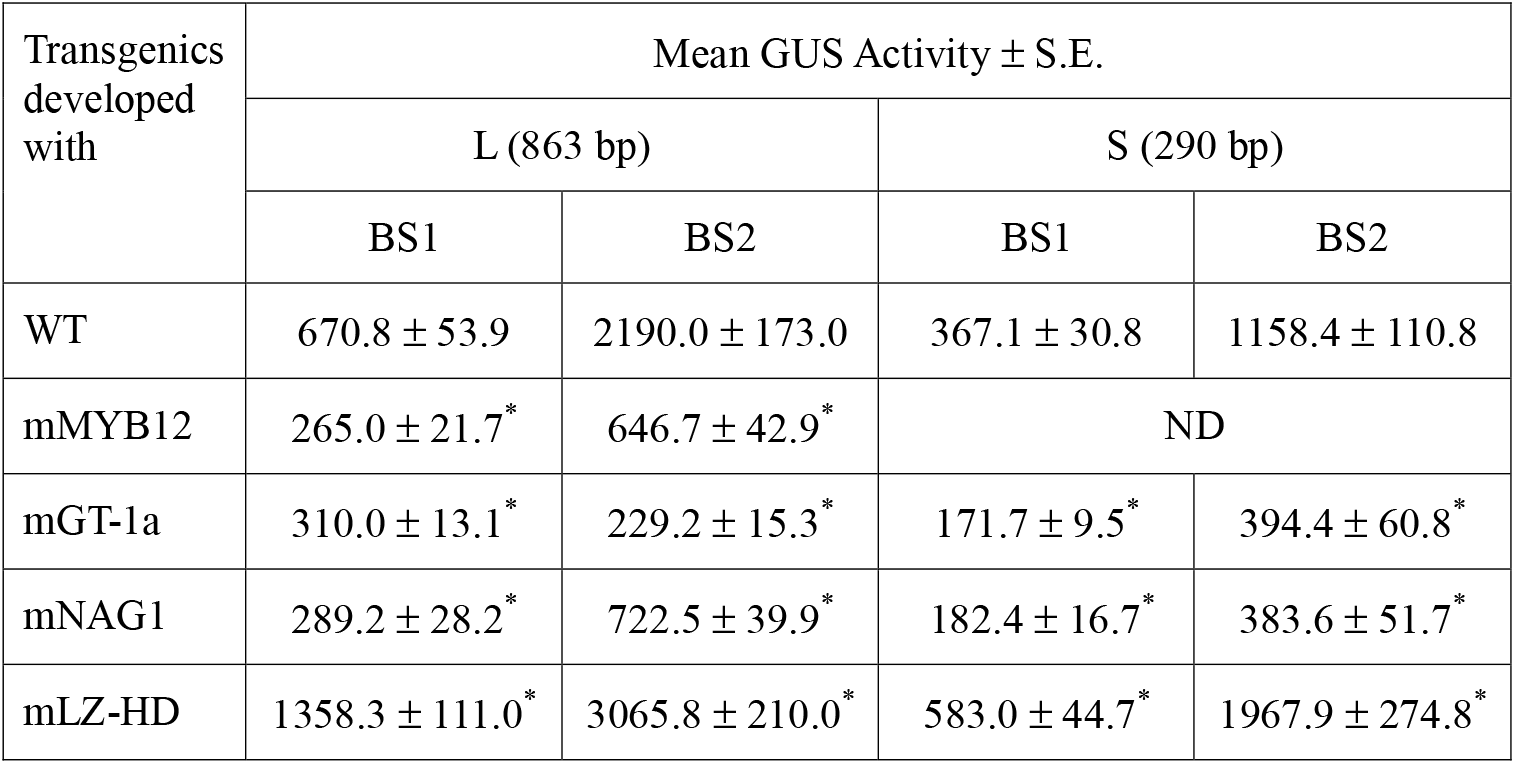
Summary of mean GUS activity observed at two different bud stages for lines developed with the different URMs. ^*^Welch’s *t*-test [SPSS version 27.0, IBM Corp, 2020] indicated that these means are significantly different from that of WT URM. Detailed analysis is presented in Supplementary Table 1. (ND – not done as the shorter version lacks a MYB 12 site.

| Transgenics developed with | Mean GUS Activity ± S.E. | | | |
| --- | --- | --- | --- | --- |
|  | L (863 bp) |  | S (290 bp) |  |
|  | BS1 | BS2 | BS1 | BS2 |
| WT | 670.8 ± 53.9 | 2190.0 ± 173.0 | 367.1 ± 30.8 | 1158.4 ± 110.8 |
| mMYB12 | 265.0 ± 21.7* | 646.7 ± 42.9* | ND | |
| mGT-1a | 310.0 ± 13.1* | 229.2 ± 15.3* | 171.7 ± 9.5* | 394.4 ± 60.8* |
| mNAG1 | 289.2 ± 28.2* | 722.5 ± 39.9* | 182.4 ± 16.7* | 383.6 ± 51.7* |
| mLZ-HD | 1358.3 ± 111.0* | 3065.8 ± 210.0* | 583.0 ± 44.7* | 1967.9 ± 274.8* |

**Fig. 3.**
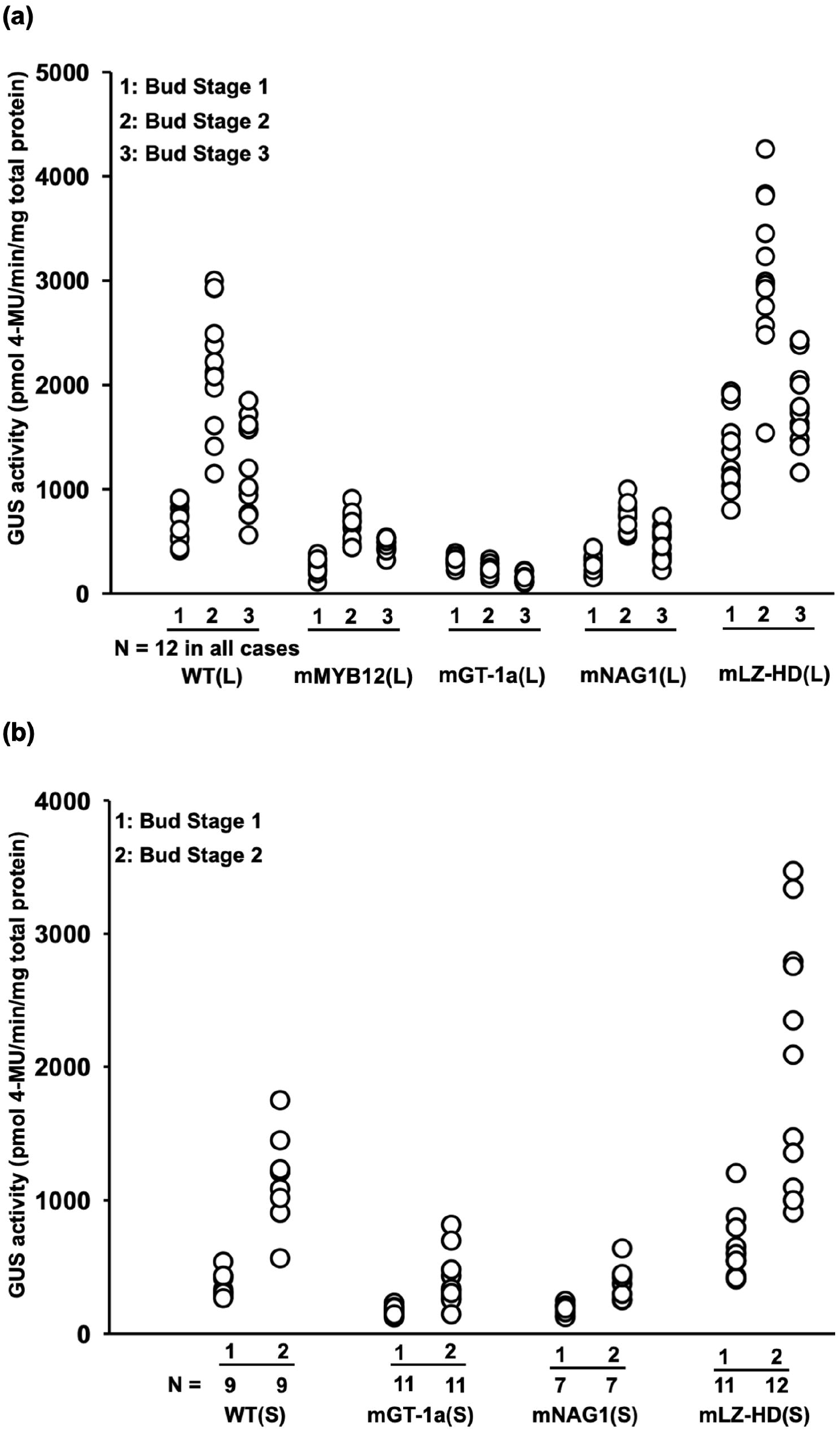
GUS activity driven by different URMs in independent transgenic lines in the background of either **(a)** −863 bp (L) or **(b)** −290 bp (S) URM.

It was observed that the activity of the wild-type URM initiated at BS1, peaked at BS2 and showed a drop at BS3. As the β-glucuronidase protein is fairly stable, the activity observed at BS3 may be a remnant of URM activity of BS2 and thus may not be a true reflection of the activity at the BS3 stage. The data clearly shows that mutant *cis*-elements for NtGT-1a and NtNAG1 led to statistically significant drop in URM activity (Table 1). Mutation of the single site for NtNAG1 located at −200 bp region led to ~50 to 57% drop in activity at BS1 and ~66 to 68% at BS2 stage, both in the longer and shorter URM background. This site is located in the 122 bp stretch of TA29 which was described as a crucial part of the TA29 promoter in the first report. This *cis*-element acts as a positive regulator for TA29 probably by binding to NtNAG1. NtNAG1 is a homolog of the *agamous* gene of Arabidopsis, which expresses in stamens and carpels, and positively regulates the TTS genes of tobacco, reported to be involved in pollen germination and tube growth (Cheung et al. 1996).

Mutation of the two sites for NtMYB12 led to a drop in URM activity ranging from ~60 to 70%. These sites though upstream to the 122 bp crucial region are within the −279 minimal promoter identified in the first report. NtMYB12 is reported to positively regulate genes involved in flavonol biosynthesis (Song et al. 2020).

Five of the six sites for NtGT-1a are located in the region upstream of −240 and one site in the putative 5 UTR. Mutation of these sites led to a drop in activity ranging from 54 to 90%, indicating their positive regulatory role. Evidence (Lam 1995) suggests NtGT-1a to be an activator. The expression profile of this gene is not established for tobacco but its homologs in *Arabidopsis* show enhanced expression in anthers.

NtLZ-HD has two sites, one at the distal end and the other at −136 bp (lying within the crucial region). Mutation of these *cis*-elements led to an increase in the activity of the URM at all stages of anther development, indicating their role as a negative regulator. The upregulation ranged from 1.4 to 2.0 folds. The TA29 URM has been reported to show expression in the meristematic regions of the stem and root in a small percentage of transgenic tobacco plants developed with WT constructs similar to those used in this study (Sharma et al. 2018). Effects of mutating the binding sites for NtLZ-HD on this ectopic expression were analysed by staining transformed callus, roots, and stems for GUS activity. It was observed (Fig. 4a) that the number of transformed events showing ectopic expression increased substantially in callus and roots of lines developed with mLZ-HD (L) constructs. The pattern of expression (Fig. 4b) remained the same as reported earlier (Sharma et al. 2018). This observation re-affirmed the role of these *cis*-elements as a negative regulator. Though the role of NtLZ-HD has not been evaluated in tobacco, its homolog in *Arabidopsis*, AtHB1, has been shown to negatively regulate the expression of miR164 (Miguel et al. 2020).

**Fig. 4a.**
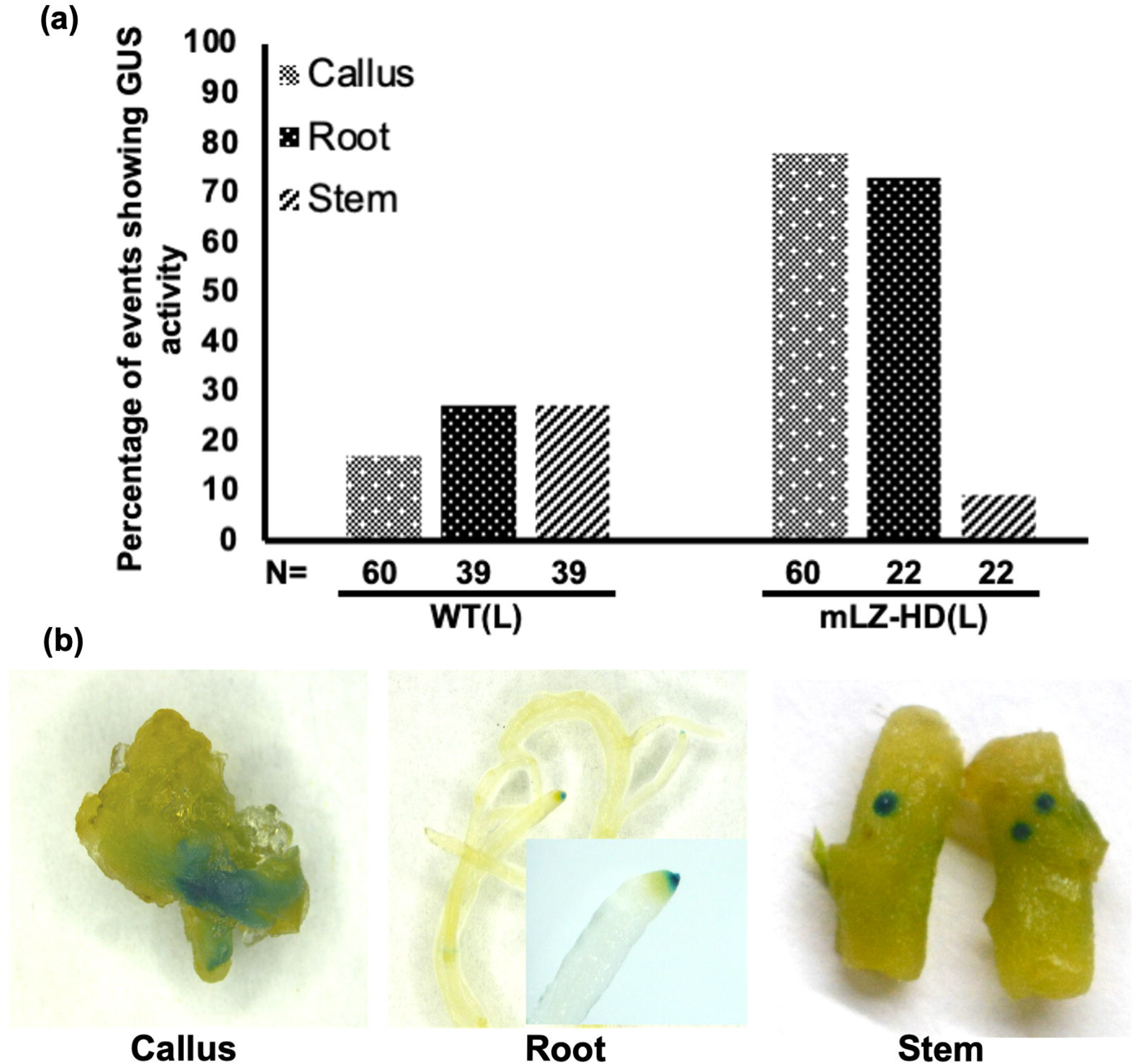
Percentage of transformed lines developed with WT(L) and mLZ-HD(L) showing GUS activity in ectopic locations i.e. callus, root and stem. No expression was observed in leaves. **(b)**. Pattern of GUS staining in the events mentioned in (a).

In summary, the current work identified *cis*-elements binding to positive (NtMYB12, NtGT-1a, NtNAG1) and negative (NtLZ-HD) regulators that are involved in regulation of the tapetum specific TA29 promoter.

## Declarations

Conflict of interest The authors declare no competing interests.

