## Supplementary Table 1 for "Tapetum specific TA29 promoter is regulated by *cis*- elements binding to positive and negative regulators"

| **Transgenics developed with** | **Number/Statistic** | **Mean GUS Activity ± S.E.** | | | | |
| --- | --- | --- | --- | --- | --- | --- |
|  |  | **L (863 bp)** | | | **S (290 bp)** | |
|  |  | **BS 1** | **BS 2** | **BS3** | **BS 1** | **BS 2** |
| WT | Range | 410.0 - 910.0 | 1150.0 - 3000.0 | 560.0 - 1850 | 266.0 - 539.5 | 564.5 - 1748 |
|  | Mean ± S.E. | 670.8 ± 53.9 | 2190.0 ± 173.0 | 1215 ± 125.7 | 367.1 ± 30.8 | 1158.4 ± 110.8 |
|  | N | 12 | 12 | 12 | 9 | 9 |
|  | W(*p*) | 0.904 (0.181) | 0.950 (0.643) | 0.924 (0.321) | 0.898 (0.243) | 0.972 (0.909) |
| mMYB12 | Range | 110.0 - 380.0 | 440.0 - 910.0 | 320.0 - 540.0 | NA | |
|  | Mean ± S.E. | 265.0 ± 21.7^***^ | 646.7 ± 42.9^***^ | 463.3 ± 18.4^***^ |  |  |
|  | N | 12 | 12 | 12 |  |  |
|  | W(*p*) | 0.962 (0.815) | 0.935 (0.437) | 0.925 (0.329) |  |  |
|  | *t*-test (df) | 6.983 (14.457) | 8.653 (12.348) | 5.918 (11.469) |  |  |
| mGT-1a | Range | 220.0 - 390.0 | 140.0 - 330.0 | 100.0 - 220.0 | 121.0 - 231.0 | 145.5 - 816.0 |
|  | Mean ± SE | 310.0 ± 13.1^***^ | 229.2 ± 15.3^***^ | 144.2 ± 11.3^***^ | 171.7 ± 9.5^***^ | 394.4 ± 60.8^***^ |
|  | N | 12 | 12 | 12 | 11 | 11 |
|  | W(*p*) | 0.989 (0.999) | 0.984 (0.994) | 0.881 (0.091) | 0.981 (0.973) | 0.874 (0.087) |
|  | *t*-test (df) | 6.501 (12.302) | 11.283 (11.172) | 8.486 (11.178) | 6.06 (9.51) | 6.046 (12.624) |
| mNAG1 | Range | 150.0 - 440.0 | 550.0 - 1000 | 220.0 - 740.0 | 123.5 - 246 | 249.5 - 638.5 |
|  | Mean ± SE | 289.2 ± 28.2^***^ | 722.5 ± 39.9^***^ | 461.7 ± 44.9^***^ | 182.4 ± 16.7^***^ | 383.6 ± 51.7^***^ |
|  | N | 12 | 12 | 12 | 7 | 7 |
|  | W(*p*) | 0.943 (0.541) | 0.934 (0.426) | 0.969 (0.904) | 0.958 (0.798) | 0.9 (0.33) |
|  | *t*-test (df) | 6.273 (16.584) | 8.26 (12.168) | 5.645 (13.764) | 5.269 (12.013) | 6.336 (11.161) |
| mLZ-HD | Range | 800.0 - 1940 | 1540 - 4260 | 1160 - 2430 | 406.5 - 873.5 | 908.5 - 3470 |
|  | Mean ± SE | 1358.3 ± 111.2^***^ | 3065.8 ± 209.5^**^ | 1785.0 ± 109.3^**^ | 583.0 ± 44.7^***^ | 1967.9 ± 274.8^*^ |
|  | N | 12 | 12 | 12 | 11 | 12 |
|  | W(*p*) | 0.926 (0.342) | 0.969 (0.902) | 0.964 (0.840) | 0.914 (0.272) | 0.887 (0.109) |
|  | *t*-test (df) | -5.562 (15.902) | -3.223 (21.245) | -3.423 (21.583) | -3.975 (16.967) | -2.732 (14.347) |

**Supplementary Table 1**: Analysis of the URM activity observed in different tobacco transgenic lines developed with 10 different constructs. N = total number of events analysed. W(*p*) indicates the Shapiro-Wilk's test statistic for normality and the probability associated with it (in parenthesis). The *t*-test (df) represent the Welch’s *t*-test values and the associated degrees of freedom (in parenthesis). The mean activity of the given URMs were compared with their respective wild type (WT) URM. ‘NA’ denotes that this site was not present in the shorter version. Asterisks after the mean ± SE values denote the significance of the *p*-values obtained after performing the *t*-test (*p ≤ 0.05, **p ≤ 0.01, ***p ≤ 0.001)
