## supplementary figure 1 for "Tapetum specific TA29 promoter is regulated by *cis*- elements binding to positive and negative regulators"

| -813 | | | | | | | | | | | | | | | | |  | | | | | | | | |  | | | | | | | | |  | | | | | | | | LZ-HD (-799 to -786) | | | | | | | | | | | | | | | | | | | | | | | | | | | | | | | | | | | | | | | | | | | | | | | | | |  | | | | | | | |  | | | | | | | | | | |
| --- | --- | --- | --- | --- | --- | --- | --- | --- | --- | --- | --- | --- | --- | --- | --- | --- | --- | --- | --- | --- | --- | --- | --- | --- | --- | --- | --- | --- | --- | --- | --- | --- | --- | --- | --- | --- | --- | --- | --- | --- | --- | --- | --- | --- | --- | --- | --- | --- | --- | --- | --- | --- | --- | --- | --- | --- | --- | --- | --- | --- | --- | --- | --- | --- | --- | --- | --- | --- | --- | --- | --- | --- | --- | --- | --- | --- | --- | --- | --- | --- | --- | --- | --- | --- | --- | --- | --- | --- | --- | --- | --- | --- | --- | --- | --- | --- | --- | --- | --- | --- | --- | --- | --- | --- | --- | --- | --- | --- | --- | --- | --- |
| T T T T T G G T T A G C G A A T G C A A T T A A T T T A G A C A T T G T G T T A | | | | | | | | | | | | | | | | | | | | | | | | | | | | | | | | | | | | | | | | | | | | | | | | | | | | | | | | | | | | | | | | | | | | | | | | | | | | | | | | | | | | | | | | | | | | | | | | | | | | | | | | | | | | | | | |
| T G T T C C A G T T A A C C G C T T C C C T G C A C T T C T T T C A A T C T A T | | | | | | | | | | | | | | | | | | | | | | | | | | | | | | | | | | | | | | | | | | | | | | | | | | | | | | | | | | | | | | | | | | | | | | | | | | | | | | | | | | | | | | | | | | | | | | | | | | | | | | | | | | | | | | | |
| C T C T C G A T A G A A A A T T G T G A T A C T T T G C G A C T T C T A T C A G | | | | | | | | | | | | | | | | | | | | | | | | | | | | | | | | | | | | | | | | | | | | | | | | | | | | | | | | | | | | | | | | | | | | | | | | | | | | | | | | | | | | | | | | | | | | | | | | | | | | | | | | | | | | | | | |
| A G G A C T T T T T G T T T T C C A T G T A A C A A T C T G T C A T T T T C G A | | | | | | | | | | | | | | | | | | | | | | | | | | | | | | | | | | | | | | | | | | | | | | | | | | | | | | | | | | | | | | | | | | | | | | | | | | | | | | | | | | | | | | | | | | | | | | | | | | | | | | | | | | | | | | | |
|  | | | | |  | | | | | | | | |  | | | | | | | | | |  | | | | | | | | |  | | | | | | | |  | | | | | |  | | | | | | | |  | | | |  | | | | | | | |  | | | | | | | |  | | | | | | | |  | | | | | | | |  | | | | | | | |  | | | | | | | | | | | | GT-1a |
| T G G G G A G A T T T G C A C A A A T A G G C T A T T T A T G T G T C C C A A T | | | | | | | | | | | | | | | | | | | | | | | | | | | | | | | | | | | | | | | | | | | | | | | | | | | | | | | | | | | | | | | | | | | | | | | | | | | | | | | | | | | | | | | | | | | | | | | | | | | | | | | | | | | | | | | |
| (2 sites; -617 to -597) | | | | | | | | | | | | | | | | | | | | | | | | | | | | | | | | | | | | | | | | | | | | | | | | | | | |  | | | | | | | | |  | | | | | | | | | | | | | | |  | | | | | | | | | | | | |  | | | | | | | | | | | | | | | |  | | | | | | |
| T T A A A T T T T A A C C C C A T G T C G A T C A G A A C T T A G C C A C G A G | | | | | | | | | | | | | | | | | | | | | | | | | | | | | | | | | | | | | | | | | | | | | | | | | | | | | | | | | | | | | | | | | | | | | | | | | | | | | | | | | | | | | | | | | | | | | | | | | | | | | | | | | | | | | | | |
| C A C C A G A A G T T T G A T G G A T A T G T G A C T T T G T C A C T A T C C G | | | | | | | | | | | | | | | | | | | | | | | | | | | | | | | | | | | | | | | | | | | | | | | | | | | | | | | | | | | | | | | | | | | | | | | | | | | | | | | | | | | | | | | | | | | | | | | | | | | | | | | | | | | | | | | |
|  | | | GT1-a (-527 to -518) | | | | | | | | | | | | | | | | | | | | | | | | | | | | | | | | | | | | | | | | | | | | | | | | | | | GT1-a (-515 to 501) | | | | | | | | | | | | | | | | | | | | | | | | | | | | | | | | | | | | | | | | | | | | | | | |  | | | | | | | |  | |
| G T T T A C T A A T C A A G A G C T A T T T T T A T T C A A A A T T G G A T A T | | | | | | | | | | | | | | | | | | | | | | | | | | | | | | | | | | | | | | | | | | | | | | | | | | | | | | | | | | | | | | | | | | | | | | | | | | | | | | | | | | | | | | | | | | | | | | | | | | | | | | | | | | | | | | | |
| C T A G C T A A G T A T A A C T G G A T A A T T T G C A T T A A C A G A T T G A | | | | | | | | | | | | | | | | | | | | | | | | | | | | | | | | | | | | | | | | | | | | | | | | | | | | | | | | | | | | | | | | | | | | | | | | | | | | | | | | | | | | | | | | | | | | | | | | | | | | | | | | | | | | | | | |
| A T A T A G T G C C A A A C A A G A A G G G A C A A T T G A C T T G T C A C T T | | | | | | | | | | | | | | | | | | | | | | | | | | | | | | | | | | | | | | | | | | | | | | | | | | | | | | | | | | | | | | | | | | | | | | | | | | | | | | | | | | | | | | | | | | | | | | | | | | | | | | | | | | | | | | | |
|  | | |  | | | | | | |  | | | | | | | | |  | | | | | |  | | | | | | |  | | | | | |  | | | | | | |  | | |  | | | | | |  | | | |  | | | |  | | | | | | | | GT1-a (-396 to -377) | | | | | | | | | | | | | | | | | | | | | | | | | | | | | | | | | | | | | | | | | |
| T A T G A A A G A T G A T T C A A A C A T G A T T T T T T A T G T A C T A A T A | | | | | | | | | | | | | | | | | | | | | | | | | | | | | | | | | | | | | | | | | | | | | | | | | | | | | | | | | | | | | | | | | | | | | | | | | | | | | | | | | | | | | | | | | | | | | | | | | | | | | | | | | | | | | | | |
| T A T A C A T C C T A C T C G A A T T A A A G C G A C A T A G G C T C G A A G T | | | | | | | | | | | | | | | | | | | | | | | | | | | | | | | | | | | | | | | | | | | | | | | | | | | | | | | | | | | | | | | | | | | | | | | | | | | | | | | | | | | | | | | | | | | | | | | | | | | | | | | | | | | | | | | |
| A T G C A C A T T T A G C A A T G T A A A T T A A A T C A G T T T T T G A A T C | | | | | | | | | | | | | | | | | | | | | | | | | | | | | | | | | | | | | | | | | | | | | | | | | | | | | | | | | | | | | | | | | | | | | | | | | | | | | | | | | | | | | | | | | | | | | | | | | | | | | | | | | | | | | | | |
|  | | | | | | | | | | |  | | | | | | | | | | | | | | | |  | | | | | | | | | | | | | | | | | (-279) | | | | | | | | | | | | | | | | | | AtMYB12 (2 sites; -272 to -260) | | | | | | | | | | | | | | | | | | | | | | | | | | | | | | | | | | | | | | | | | | | | | | | | | |
| A A G C T A A A A G C A G A C T T G C A T A A G G T G G G T G G C T G G A C T A | | | | | | | | | | | | | | | | | | | | | | | | | | | | | | | | | | | | | | | | | | | | | | | | | | | | | | | | | | | | | | | | | | | | | | | | | | | | | | | | | | | | | | | | | | | | | | | | | | | | | | | | | | | | | | | |
|  | | | | | | |  | | | | | | | |  | | | | | | | |  | | | | | | |  | | | | | | | (-240) | | | | | | | | | | | | | | | | | | | | | | | | | | | | | | | | | | | |  | | | | | | | | | | | | | | | | | | | | | | | | |  | | | | | | | | | | NAG1 | | | |
| G A A T A A A C A T C T T C T C T A G C A C A G C T T C A T A A T G T A A T T T | | | | | | | | | | | | | | | | | | | | | | | | | | | | | | | | | | | | | | | | | | | | | | | | | | | | | | | | | | | | | | | | | | | | | | | | | | | | | | | | | | | | | | | | | | | | | | | | | | | | | | | | | | | | | | | |
| (-217 to -200) | | | | | | | | | | | | | | | | | | | | | | | | | | |  | | | | | | | | | | | | | | | | |  | | | | | | | |  | | | | | | | | | |  | | | | | | | | | | | | | | |  | | | | | | | | | | | | |  | | | | | | | | | | | | | | | |  | | | | | |
| C C A T A A C T G A A A T C A G G G T G A G A C A A A A T T T T G G T A C T T T | | | | | | | | | | | | | | | | | | | | | | | | | | | | | | | | | | | | | | | | | | | | | | | | | | | | | | | | | | | | | | | | | | | | | | | | | | | | | | | | | | | | | | | | | | | | | | | | | | | | | | | | | | | | | | | |
|  | | | |  | | | | | | | | |  | | | | | | | |  | | | | | | | | | |  | | | | | | | |  | | | | | | | | | | | | | | | | | | | | | | | |  | | | | | | | | | LZHD (-148 to -136) | | | | | | | | | | | | | | | | | | | | | | | | | | | | | | | | | | | | | | | |
| T T C C T C A C A C T A A G T C C A T G T T T G C A A C A A A T T A A T A C A T | | | | | | | | | | | | | | | | | | | | | | | | | | | | | | | | | | | | | | | | | | | | | | | | | | | | | | | | | | | | | | | | | | | | | | | | | | | | | | | | | | | | | | | | | | | | | | | | | | | | | | | | | | | | | | | |
| G A A A C C T T A A T G T T A C C C T C A G A T T A G C C T G C T A C T C C C C | | | | | | | | | | | | | | | | | | | | | | | | | | | | | | | | | | | | | | | | | | | | | | | | | | | | | | | | | | | | | | | | | | | | | | | | | | | | | | | | | | | | | | | | | | | | | | | | | | | | | | | | | | | | | | | |
| A T T T T C C T C G A A A T G C T C C A A C A A A A G T T A G T T T T G C A A G | | | | | | | | | | | | | | | | | | | | | | | | | | | | | | | | | | | | | | | | | | | | | | | | | | | | | | | | | | | | | | | | | | | | | | | | | | | | | | | | | | | | | | | | | | | | | | | | | | | | | | | | | | | | | | | |
| T T G T T G T G T A T G T C T T G T G C T C T A T A T A T G C C C T T G T G G T | | | | | | | | | | | | | | | | | | | | | | | | | | | | | | | | | | | | | | | | | | | | | | | | | | | | | | | | | | | | | | | | | | | | | | | | | | | | | | | | | | | | | | | | | | | | | | | | | | | | | | | | | | | | | | | |
|  | |  | | | | | |  | | | | | | | |  | | | | |  | | | | | | | |  | | | | | | |  | | | | TSS(+1) | | | | | | | | | | | | | | | | | | | | | | | | | |  | | | | |  | | | | | | |  | | | | | |  | | |  | | | | | | |  | | | | | |  | | | | | | |  | | | | |
| G C A A G T G T A A C A G T A C A A C A T C A T C A C T C A A A T C A A A G T | | | | | | | | | | | | | | | | | | | | | | | | | | | | | | | | | | | | | | | | | | | | | | | | | | | | | | | | | | | | | | | | | | | | | | | | | | | | | | | | | | | | | | | | | | | | | | | | | | | | | | | | | | | | | | | |
| GT1-a (+27 to +37) (+50) | | | | | | | | | | | | | | | | | | | | | | | | | | | | | | | | | | | | | | | | | | | | | | | | | | | | | | | | | | | | | | | | | | | | | | | | | | | | | | | |  | | | | |  | | | | | | | | | |  | | | | | | | | |  | | | | | | | |
| T T T T A C T T A A A G A A A T T A G C T A C C A T G | | | | | | | | | | | | | | | | | | | | | | | | | | | | | | | | | | | | | | | | | | | | | | | | | | | | | | | | | | | | | | | | | | | | | | | | | | | | | | | | | | | | | | | | | | | | | | | | | | | | | | | | | | | | | | | |

Supplementary Figure 1: The sequence of TA29 URM (863 bp) is presented. The URM consists of a 50 bp 5´UTR. The putative transcription start site (TSS) has been indicated (	). The identified putative *cis*-elements binding to different transcription factors has been indicated. The (	) indicates distal end of the 290 bp TA29 URM used in the study The crucial region identified by Koltunow et al (1990) has been underlined. Translation start site (ATG) is marked in larger font.
